# hadge: a comprehensive pipeline for donor deconvolution in single cell

**DOI:** 10.1101/2023.07.23.550061

**Authors:** Fabiola Curion, Xichen Wu, Lukas Heumos, Mariana Gonzales, Lennard Halle, Melissa Grant-Peters, Charlotte Rich-Griffin, Hing-Yuen Yeung, Calliope A. Dendrou, Herbert B. Schiller, Fabian J. Theis

**Author notes:** These authors contributed equally (Fabiola Curion, Xichen Wu, Lukas Heumos).

## Abstract

Single cell multiplexing techniques (cell hashing and genetic multiplexing) allow to combine multiple samples, thereby optimizing sample processing and reducing batch effects. Cell hashing conjugates antibody-tags or chemical-oligonucleotides to cell membranes, while genetic multiplexing allows to mix genetically diverse samples and relies on aggregation of RNA reads at known genomic coordinates. We developed hadge (**ha**shing **d**econvolution combined with **ge**notype information), a Nextflow pipeline that combines 12 methods to perform both hashing- and genotype-based deconvolution. We propose a joint deconvolution strategy combining the best performing methods and we demonstrate how this approach leads to recovery of previously discarded cells in a nuclei hashing of fresh-frozen brain tissue.

## Introduction

Single cell RNA sequencing (scRNA–seq) technologies have unlocked unprecedented resolution to discover complex mechanisms of health and disease in human biology^1^. Droplet-based methods, which encapsulate aqueous cells into oil constituting a micro-chamber for lysis and retrotranscription of the RNA of individual cells, have made single cell sequencing more accessible and dramatically increased the throughput of single cells from individual samples^2^. The cDNA produced in these reactions is uniquely barcoded for each droplet, such that the retrieval of the barcodes allows the association of sequencing readouts to individual cells.

Despite the considerable strides made in cellular profiling methods, including the continuous reduction in costs, the acquisition of fresh tissue samples for single cell research remains a challenge. This is particularly true for specimens from biobanks, where such samples may not be readily available. Furthermore, the need to profile multiple patients and tissues to increase statistical power compounds the complexity and cost of single cell research. These challenges emphasize the pressing need for the development of new technological approaches to enable efficient and cost-effective analysis of diverse cellular populations.

In recent years, methods have emerged that allow multiplexing of single cells from individual samples by pooling samples with distinct genotypes together into a single experiment. This reduces experimental cost and increases the throughput of large, multi-sample experiments^3^. These methods have found wide applicability and have already allowed profiling of thousands of cells from hundreds or thousands of human samples, unlocking the statistical power to carry out population studies with single cell techniques^4, 5^.

To date, there are two major ways to generate and deconvolve (also sometimes referred to as demultiplexing) a mixture of samples: “cell hashing” and “genotype-based multiplexing”. Cell hashing is a sample processing technique that requires processing individual samples to “tag” the membrane of the cell or the nuclei with unique oligonucleotide barcodes. One option is to use oligonucleotide-labeled antibodies that target proteins ubiquitously expressed on the cell or nuclei surface. Another option is to chemically conjugate oligonucleotides directly to the membrane constituents, for example by hybridization of a lipid-modified oligonucleotide (LMO) to the hydrophobic cell membrane, a technique called “lipid tagging”^6^, or by chemical ligation of the oligonucleotide to exposed N-Hydroxysuccinimide-reactive amines, a technique called “chemical barcoding”^7–9^. After cell tagging, the cells are washed or the reaction is quenched, and the samples can be safely mixed and processed following the standard library preparation procedure. Two libraries are generated after this process, one for the scRNA and one for the hashing oligos (HTO), which are independently sequenced to produce each a single cell count matrix, one for the RNA library and one for the HTO library. The hashtag counts are then bioinformatically processed to deconvolve the cell’s source sample (**Fig. 1A**). However, cell-tagging approaches may not be appropriate if the starting cell numbers are a limiting factor, as these methods require washing steps that may result in cell-number loss. Furthermore, different issues can impair the quality of a hashing experiment, and therefore decrease the final number of uniquely identified cells. Antibodies or free oligos can persist in suspensions if an adequate number of washes is not performed, or can attach to debris from membrane lysis in fixed samples^10^.

**Figure 1.**
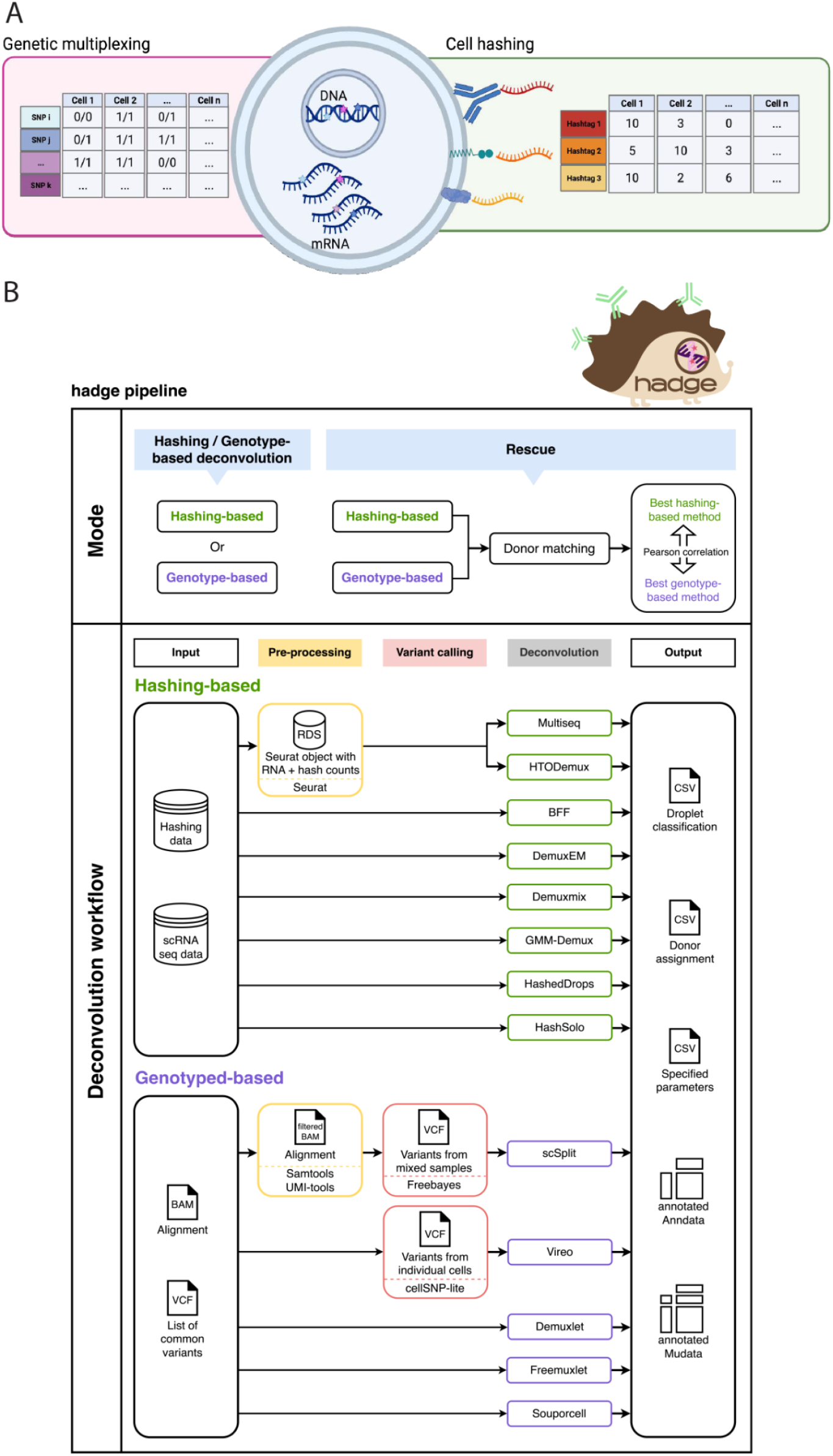
Overview of donor deconvolution and the hadge pipeline. (A) Schematic example of the cellular components leveraged by single cell multiplexing experiments. The output data (i.e. hashtag cell-counts or SNP calling by cell) is the input to the hadge pipeline. (B) hadge implements 10 methods across two sub-pipelines, hashing-based and genotype-based deconvolution, which can be run independently, in parallel or in rescue mode. In rescue mode, the pipeline offers the option to refine hashing results with genotype-based deconvolution methods to rescue failed hashing experiments in the donor matching process. It compares the concordance in donor identification between hashing and genotype-based methods and identifies the best pair of two strategies based on Pearson correlation

Genotyped-based deconvolution instead leverages the unique genetic composition of individual samples to guarantee that the final cell mixture can be deconvolved. Users can genotype the individual samples through single nucleotide polymorphism (SNP) arrays or bulk RNA-seq followed by variant calling, and then aggregate the expression values at these genomic positions to deconvolute the samples identities. The same process can be conducted without genotype of origin (“genotype-free”) by piling up the mixture scRNA onto a genomic reference from unmatched donors, for example the 1000 genome project genotypes^11^. The result of this approach is a table of SNP assignment to cells that can be used to computationally infer the donors (**Fig. 1A**). One limitation of this approach is the need to produce additional data to genotype the individual donors in order to correctly assign the cell mixtures.

Cell hashing and genotype-based multiplexing techniques have been applied in a variety of conditions and tissues, with considerably variable performance depending on the efficiency of the protocols and the quality of the input samples^9, 12–14^.

To date, at least nine hashing and five genotype-based deconvolution methods have been developed, each with unique strengths and weaknesses^6, 10, 15–21^. Although workflows for hashing-based deconvolution and genotype-based deconvolution exist^22, 23^, no study has combined all these tools in a single comprehensive pipeline, such that both hashing and genotype deconvolution pipelines can be run in parallel on multiple samples. Moreover, the joint call of hashing and genetic deconvolution methods has been shown to be beneficial for cell recovery rate and calling accuracy^24^. These investigations have been limited to the combination of two tools instead of computationally testing the best combination of demultiplexing methods, therefore neglecting the utility of other widely used tools. As such, there is a critical need for a unified pipeline that integrates the strengths of multiple donor deconvolution tools.

Here we present the hadge (**ha**shing **d**econvolution combined with **ge**netic information) pipeline. Our Nextflow^25^ based pipeline enables deconvolving samples of both hashing and genetic multiplexing experiments either independently or simultaneously. hadge allows for the automatic determination of the best combination of hashing and SNP- based donor deconvolution tools. Moreover, hadge provides a rescue mode to run both genetic and hashing approaches jointly to rescue problematic hashing experiments in cases where donors are genetically distinct. We demonstrate our pipeline using a single nuclei hashing experiment of fresh frozen multiple sclerosis (MS) brain tissue and show that joint deconvolution allows us to rescue high quality cells that would have been otherwise discarded.

## Results

### The hadge pipeline

hadge offers a user-friendly, zero-config solution for analyzing single cell sequencing data at scale (**Fig. 1B**). Our pipeline takes advantage of Nextflow’s cloud-computing capabilities, enabling efficient use of cloud resources to accelerate analysis of large datasets. Furthermore, Nextflow’s built-in containerization functionality simplifies deployment, providing a more reliable and reproducible analysis environment. The hadge pipeline consists of 10 deconvolution tools, including five genetics-based tools (Demuxlet^17^, Freemuxlet^26^, Vireo^21^, scSplit ^19^, and Souporcell^20^), eight hashing-based tools (HTODemux^27^, Multiseq^6^, HashedDrops^16^, Demuxem^10^, gmm-demux^28^, BFF ^23^, demuxmix^29^ and Hashsolo^18^), and one doublet-detection method (Solo^18^). Furthermore, for methods that require additional preprocessing, such as normalization of the HTO counts or variant calling, the hadge pipeline includes a preprocessing step before the genotype-based deconvolution algorithm is applied.

The hadge pipeline has three modes: “genetic”, “hashing” and “rescue”. In the *genetic* or *hashing* mode, the pipeline runs either the genotype- or hashing-based deconvolution pipeline allowing for choice of methods and customization of input parameters. Each of these pipelines can be run in parallel across multiple samples, reducing the time and effort required for deconvolution. Finally, in the *rescue* mode, hadge allows jointly deconvolving hashing experiments with genotype-based deconvolution tools, with the option to recover the cells from failed hashing. Since the genotype-based deconvolution tools are run in a genotype-free mode, they output their assignment in the form of anonymous donors. After conversion of the cell deconvolution into a binary matrix with rows representing cell barcodes and columns representing the assigned donors or hashtags, donor genotypes are matched with hashtags by measuring the concordance of two methods in assigning the droplets, computing pairwise Pearson correlation to determine the optimal match. hadge then generates a new assignment of the cells based on this optimal match between hashing and genotype-based deconvolution to uncover the true donor identity of the cells effectively rescuing cells from failed hashing with a valid genotyped-based deconvolution assignment. Finally, hadge outputs the results of the donor deconvolution for all combinations of methods and hyperparameters tested, both as a separate tabular format, and as cell metadata in either Anndata^30^ or MuData^31^ objects, depending on the users’ choice.

### Hashing-based methods performance greatly varies with noisy HTO libraries

We applied the hadge pipeline to a hashing dataset of single nuclei sequencing collected from post-mortem brain tissue from multiple sclerosis donors^32^. The hashing count matrix of this dataset presented a high background noise from non-specific antibody binding (**Fig. 2A**). We ran both hashing and genotyped-based deconvolution workflows with the aim of assessing the performance of the two types of approaches. We observed inconsistent hashtags counts (**Fig. 2A, 2B** and **Supp. Fig. 1**). Specifically, hashtag 453 showed a high overall expression, while hashtags 454 and 455 were expressed in relatively low levels (**Fig. 2A and Supp. Fig. 1, 6**). Due to the variable readout of the hashing oligos, the sample-assignment of the hashing-based methods was not consistent. The number of detected singlets varied greatly between different methods (**Fig. 2C and Supp. Fig. 1-2, 7**). While Hashsolo classified almost every droplet as a singlet, HashedDrops detected only 32 singlets among 4048 non-empty droplets. Demuxem and Multiseq exhibited similar performance, assigning about 1,800 singlets. Single nuclei hashing experiments may be more affected by unwanted background noise when compared to single cell hashing, because the broken cell membranes cause debris which can cause non-specific antibody-binding^10^. However, the expression profiles of these cells are still of good quality which allows genotypes to be called from the RNA reads. Compared to hashing, genotype-based deconvolution methods performed more consistently and identified significantly more singlets (**Fig. 2D and Supp. Fig. 3, 8-9)**. Each tool classified over 90% of the droplets as singlets, and there was consistent agreement between all tools for 3914 singlets (**Fig. 3D**). However, scSplit identified 296 droplets as doublets, which were consistently identified as singlets by three other methods. Due to the high consistency among Vireo, Freemuxlet and Souporcell, and available benchmarks showcasing its favourable performance compared to the other tools ^22^ we decided to use Vireo as a baseline for genotype-based deconvolution methods.

**Figure 2.**
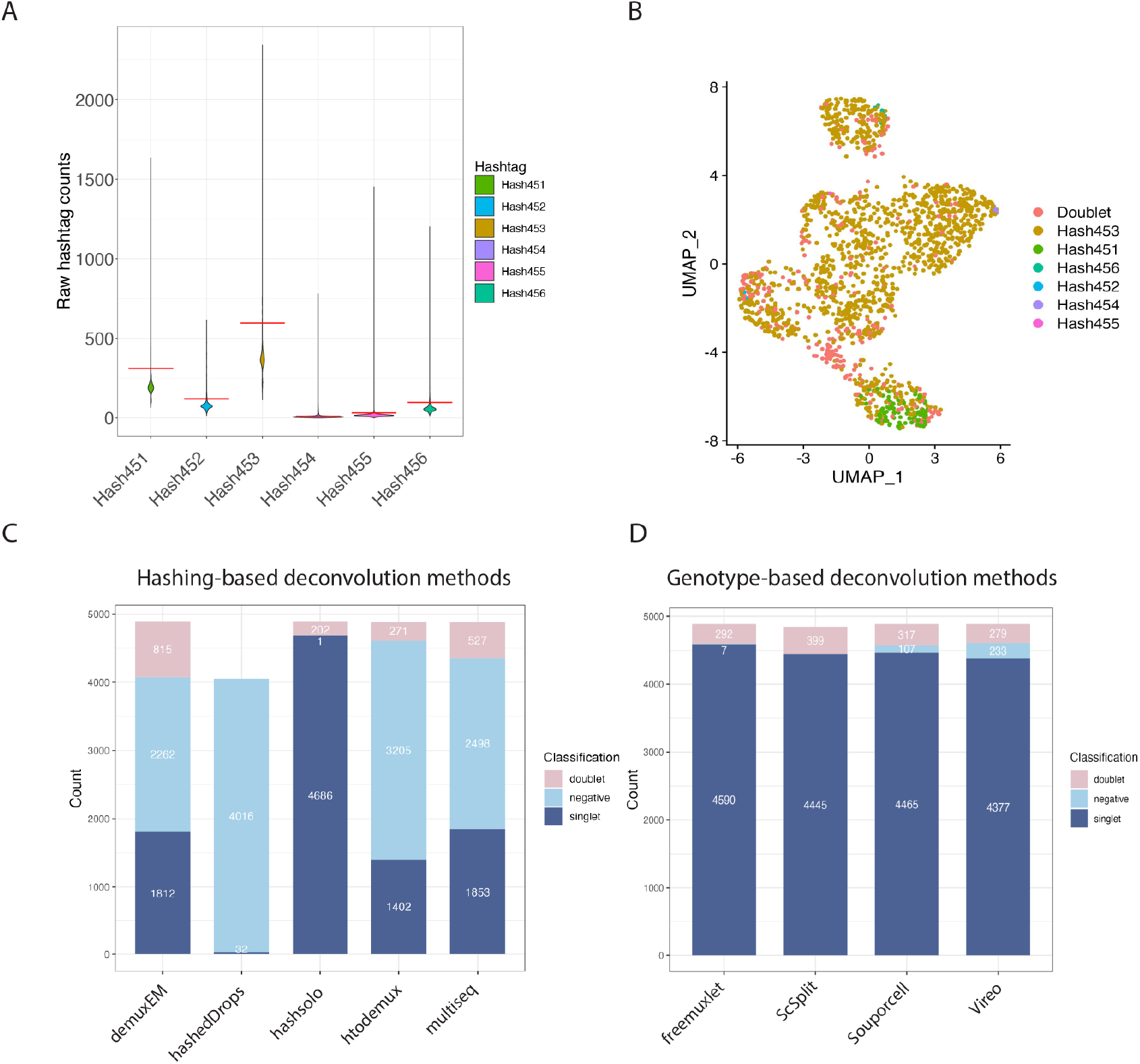
Comparison of the performance of donor deconvolution methods. (A) The violin plot of raw HTO counts shows a high counts levels of Hashtag 453 in cells with noisy or undetectable expression of the other HTOs. (B) t-SNE plot of normalized HTO counts colored by HTODemux assignment shows poor separation of the cells based on hashtags, with most droplets assigned to Hashtag 453. (C) Hashing-based deconvolution methods show inconsistent assignment of cells, reported as the different proportions of cells identified as one of either singlet, negative or doublet. (D) Genetic deconvolution tools show a more consistent assignment of the cell mixture to singlets, doublets and negatives.

**Figure 3.**
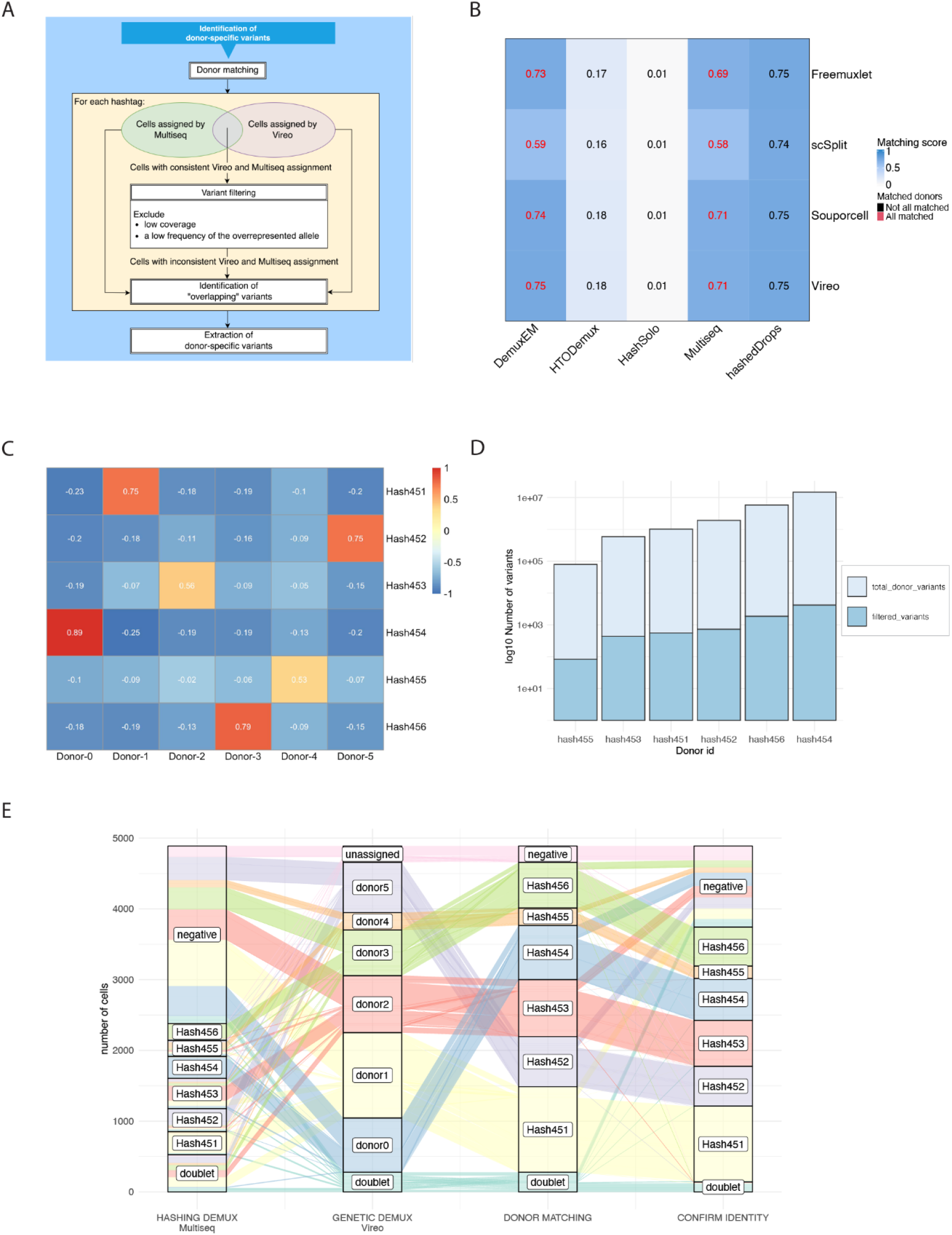
Joint deconvolution recovers high quality cells. (A) Overview of extracting cell-variants from common SNPs in the population based on the assignment of Multiseq and Vireo. (B) Heatmap summarizing the donor matching result shows that DemuxEM and Multiseq are in high concordance with all genotype-based deconvolution methods, where all the donors are matched with a high matching score. (C) Correlation heatmap of donor identification between Vireo and Multiseq. (D) Number of donor-specific variants before and after filtering that are used as input for optional donor refinement (E) Alluvial plot summarizing the subsequent changes in annotation of the cells as they are passed through the four steps of the hadge pipeline. Each column reports the number of cells deconvolved to individual donors at each step. The first two columns show the results of hashing- and genotype-based deconvolution, determined by Multiseq and Vireo respectively. The third column refers to the result of donor matching, where each anonymous donor genotyped by Vireo is matched to a hashtag. The fourth column is the assignment of Vireo with reconstructed donor genotypes.

### Joint deconvolution recovers cells with low quality hashing data

hadge additionally aims to determine the optimal combination of hashing- and genotype-based deconvolution methods and to rescue cells whose hashing quality was low or whose hashes were missing. Since donor-specific reference genotypes are not available, all genotype-based deconvolution tools output their assignment in the form of anonymous donors, which complicates the assessment of concordance between methods from two different strategies. To assign the recovered genetic singlets to the original donors, we developed a matching score based on Pearson correlation. We perform donor matching between hashing, where the identity of the donors is known, and genetic classification, by computing the Pearson correlation on the binarized classification vectors (Methods). The degree of consistency in donor identification between any two methods is evaluated by the matching score, which is calculated by summing the Pearson correlation scores of the paired hashtag and anonymous donor cluster. Based on the observed high matching score and the successful matching of all anonymous donors (**Supp. Fig. 4-5, 10**), two hashing demultiplexing methods performed best compared to Vireo, namely Multiseq and Demuxem (**Fig. 3B**). Here, we decided to use the joint demultiplexing of Multiseq and Vireo to showcase the *rescue* mode (**Fig. 3A)**. For every anonymous donor recovered by Vireo there was only one hashtag with high Pearson correlation, with scores ranging from 0.53 to 0.89 (**Fig. 3C**). This enabled us to map donor IDs to hashtags to reveal the true identity of the genotype clusters, and extend the classification to those cells whose hashing was undetectable (negatives), rescuing 89.7% of the original negatives (**Fig. 3C, 3E**). In hadge, Vireo is implemented to rely on the file outputs generated by cellSNP. The pipeline offers an optional process to refine the cell deconvolution by extracting cell-variants in order to reconstruct the donor genotypes from the common SNPs in the populations. Variants with low coverage (allele depth <10) or a low frequency of the overrepresented allele (frequency < 0.1) were excluded (**Fig. 3D**), revealing 7866 variants that were unique to each donor. Using this refining approach, the number of rescued negatives decreases to 69.7%, but with 97.6% consistent donor assignment between the rescued and the refined assignments (**Fig. 3E**), suggesting that these variants were probably crucial in distinguishing a donor cluster from others during deconvolution.

**Figure 4.**
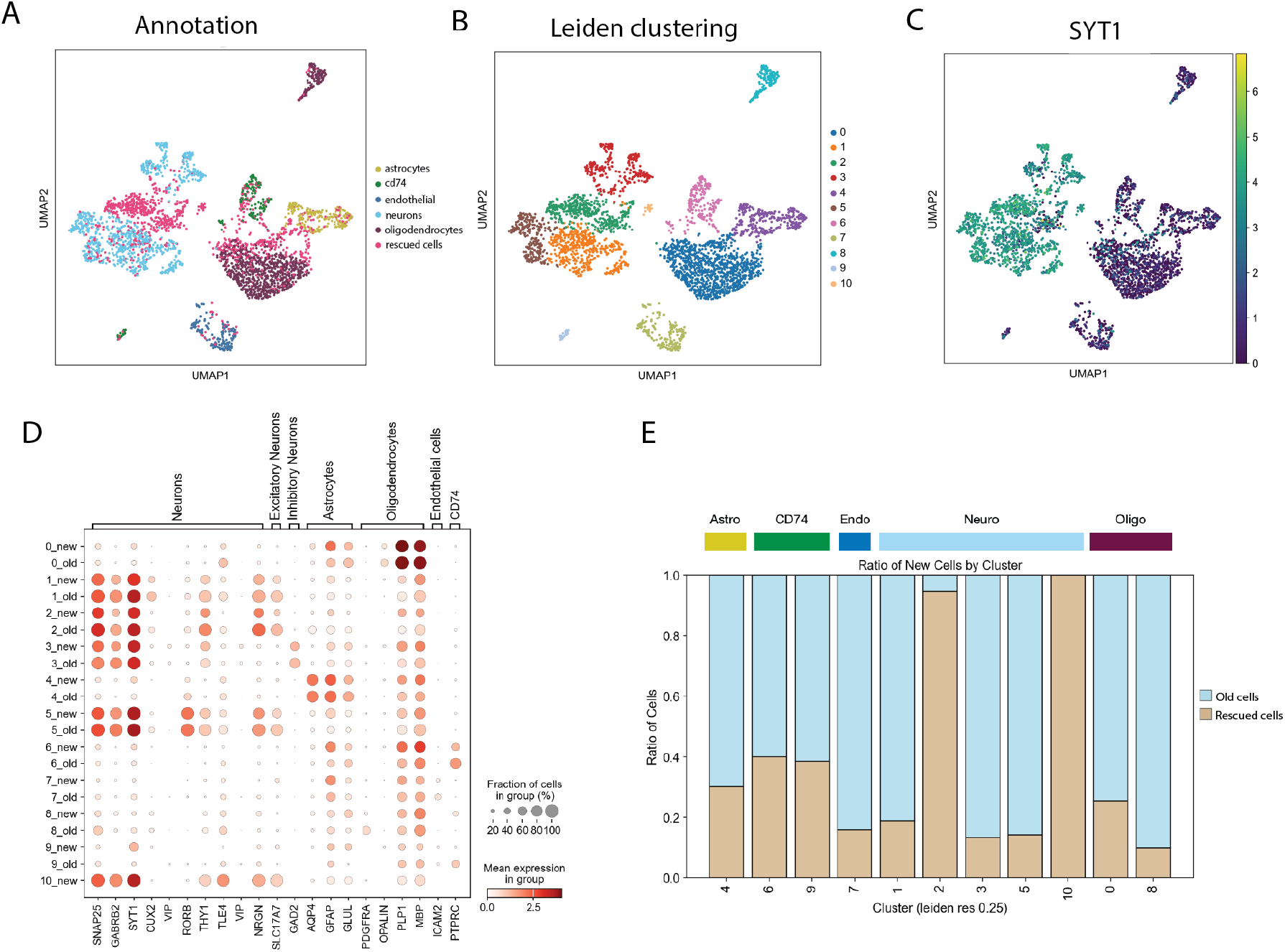
Recovered cells recapitulate known cell types (A) UMAP of the single cell gene expression data with old and rescued cells. (B) Leiden clustering of the dataset with old and rescued cells. (C) SYT1 expression defines rescued cells as a new cluster of neurons. (D) Dotplot of a selection of marker genes shows concordant expression of markers in old and rescued cells. (E) Barplot showing the cluster-composition in old and rescued cells, with two neuronal clusters enriched for rescued cells. Colors on top of barplot identify the cell annotation from (A).

**Figure 5.**
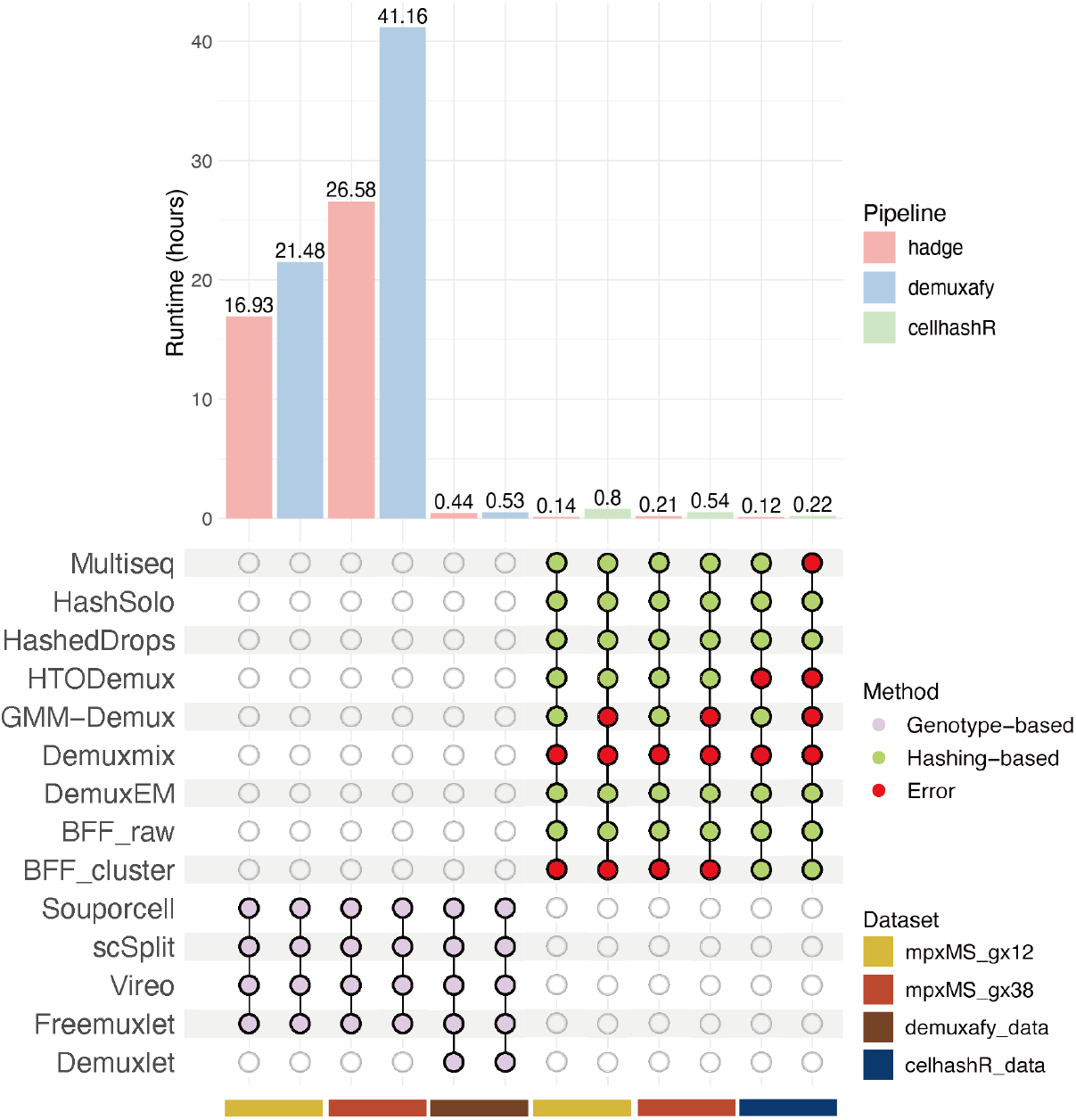
Benchmarking performance. Hadge genetic and hashing demultiplexing pipelines were benchmarked against demuxafy and cellhashR. The benchmark was performed on three samples for each pairwise comparison, for a total of four samples. (mpxMS_gx12, mpxMS_gx38, demuxafy_data, cellhashR_data). The individual pipelines were run on a linux server requesting 32 CPU cores and 160GB RAM each.

### Recovered cells recapitulate known cell types

To investigate whether the cells that are rescued are of good quality and biologically relevant, we reanalyzed the sample including the recovered cells. We first merged the already existing annotation of the cells with the deconvolution-information obtained from the hadge pipeline. We then removed cells based on gene content, mitochondrial percentage and doublet rates (Methods), reproducing the quality control performed in the original study but with a more stringent doublet detection threshold. With this approach, we retained 3,208 cells, rescuing 952 cells that were excluded in the original study. We then embedded the cells using UMAP and calculated Leiden clustering. Most of the rescued cells were distributed across existing clusters, with comparable marker expression between the old and new cells (**Fig. 4A-B,D, Supp. Fig. 11**). Intriguingly though, the percentage of rescued cells per cluster varied. While most of the clusters consisted predominantly of previously annotated cells mixing with a smaller part of rescued cells, two clusters were composed of more than half or even one hundred percent rescued cells (**Fig. 4E**). While the smaller one of these, consisting solely of rescued cells, had an almost exclusively high expression of the marker *HTR2C*, we found the gene marker expression of e.g. *SYT1*, *SLC17A7* and low *GAD2* to be consistent with a neuronal profile with excitatory and non-inhibitory properties in both clusters. Reassuringly, the latter marker expression was in accordance with that of known neuronal clusters (**Fig. 4C,D, Supp. Fig. 11)**.

### Benchmarking hadge’s performance

To demonstrate the functionality of our proposed pipeline, we benchmarked its performance against 2 existing pipelines, demuxafy and cellHashR (**Table 1)** We submitted each pipeline on a Linux server requesting for each 32 CPU cores and 160GB of RAM memory. In all benchmarks, hadge showed superior performance with respect to the optimization of computational resources and runtime. (**Fig. 5**) Both *hadge-genetic* and *demuxafy* ran all the included methods successfully for the two mpxMS samples and the additional dataset tested. For the the hashing deconvolution of the mpxMS data, for both *hadge-hashing* and *cellhashr*, some methods were successfully ran but failed to deconvolve the cells (*bff_cluster, bff_raw),* while one method failed at the startup for both pipelines and as standalone method (*demuxmix*). Notably, despite running successfully *gmm_demux* within *hadge-hashing* or when called outside the pipeline, we were not able to run cellhashR’s *gmm-demux* module.

**Table 1.** Comparison of donor deconvolution pipelines. Both “Pre-processing tools” and “variant calling tools” feature showcases the supported tools in the pipeline. “Concatenating” refers to functionality to concatenate hashing-based and genotype-based deconvolution methods. “Combining Results” refers to the functionality that allows the merging of results from multiple methods into a single data frame during a single run, based on the users’ choice. (+) The pipeline supports the mentioned functionality. (-) The pipeline doesn’t support the mentioned functionality. (*) The software is required as part of additional preprocessing outside of the pipeline.

|  | Demuxafy | cellHashR | HTOreader | hadge |
| --- | --- | --- | --- | --- |
| Framework | Singularity | R | R | Nextflow |
| Available genotype-based methods | 5 | - | Souporcell | 5 |
| Available hashing-based methods | - | 7 | HTOreader | 8 |
| Doublet detection methods | 7 | - | - | 1 |
| Concatenating | - | - | -(*) | + |
| Parallelized | - | - | - | + |
| Pre-processing tools | Samtools (*) | ProcessCount<br>Matrix,<br>PlotNormaliz<br>ationQC | HTOClassification | Samtools |
| Variant calling tools | Freebayes(*),<br>cellSNP-lite (*) | (not relevant for<br>hashing-based) | (not relevant for<br>hashing-based) | Freebayes,<br>cellSNP-lite |
| Associating clusters and donors | Only though<br>reference SNP<br>genotypes | (not relevant for<br>hashing-based) | From hashtags to<br>donors based on<br>confusion matrix | From hashtags to<br>donors based on<br>Pearson<br>correlation |
| Combining results | + | + | - | + |
| scverse compatibility | - | - | - | + |

## Discussion

Single cell multiplexing techniques enhance sample throughput, reduce costs, minimize technical variation, and improve cell type identification in single cell genomics studies by increasing the number of samples and therefore reducing the gene expression variation associated with single cell RNA sequencing. Some of the techniques for generating multiplexed single cell mixtures require additional processing steps, which can introduce technical noise and result in a low yield of usable data. Furthermore, computational donor deconvolution errors can occur due to technical noise or experimental artifacts, leading to misidentification of cells or barcodes.

We developed hadge, a comprehensive pipeline for donor deconvolution experiments generated with both genetic and hashing multiplexing methods. hadge is the only pipeline capable of processing both types of data inputs allowing for fine tuning of deconvolution experiments. To ensure confident identification of cell mixtures, hadge enables complete customization of input hyperparameters and selection of methods, and offers a host of diagnostic plots and statistics to compare results between methods. Furthermore, hadge performs joint genotype- and hashing-based deconvolution of cell mixtures generated from genetically diverse inputs to enable users to retrieve only confidently assigned singlets. This functionality is particularly relevant for experiments where cell-tagging data quality may be compromised by technical noise, tissue-specific variability, or variability in reagent performance. In these experiments, genotype-free deconvolution followed by donor matching can increase the number of good quality singlets which can be further investigated for biological signatures. Another recent work ^24^ proposed joint deconvolution to increase the confidence in called singlets, but offers limited options to customize the selection of tools or parameters to run the joint deconvolution step (**Table 1**). Given the importance of retaining only correctly assigned cells for downstream tasks, such as cell annotation and differential expression between multiplexed conditions, joint deconvolution is a necessary step for experiments threatened by suboptimal cell-tagging libraries. Existing strategies generally only retain the union of singlets called by two methods ^32^. Instead, hadge allows both automated matching of the best hashing and genotype-based deconvolution tools based on the optimal concordance between methods, or the selection of individual methods for each workflow, ensuring an additional level of control over the joint deconvolution step. To guarantee that the joint deconvolution is retaining only confidently donor-assigned singlets, we developed an additional component that allows the generation of donor genotypes from recovered single cell variants, which are then used as input for a new round of deconvolution. One limitation of this approach is that, by reducing the number of input variants to include only donor-specific variants, the read coverage in the already shallow-depth single cell data may decrease at individual genetic variants, resulting in a higher number of cells discarded as negatives. Nevertheless, in the data presented here, only fifteen (0.03%) of the total cells are misclassified into a different donor at this step, suggesting the relevance of the selected genetic variants.

Other pipelines have been proposed to benchmark either genotype-based^22^ or hashing-based deconvolution^13, 23^ individually (**Table 1**). However, some deconvolution tools do not integrate well with downstream analysis pipelines, making it difficult to perform integrated analyses across multiple samples or experiments. hadge seamlessly integrates within the scverse^33^ ecosystem, and its outputs can be processed with existing pipelines for automated single cell analysis^34^, minimizing the friction between preprocessing and data analysis steps and ensuring quality and reproducibility of results.

In conclusion, hadge is a powerful and flexible pipeline that addresses the challenges associated with all commercially available single cell multiplexing techniques in genomics studies. By allowing customization of input parameters, selection of methods, and joint deconvolution, hadge ensures confident identification of cell mixtures and retrieval of high-quality singlets. Its integration with existing analysis pipelines and compatibility with the scverse ecosystem further streamlines the data processing and analysis workflow, promoting reproducibility and enabling integrated analyses across multiple samples and experiments. With its comprehensive features and robust performance, hadge is poised to greatly enhance the accuracy and efficiency of single cell genomics research.

## Methods

### Implementation of the hadge pipeline

The hadge pipeline, implemented in Nextflow, provides hashing- and genotypes-based deconvolution workflows. Both workflows support the execution of multiple methods simultaneously.

#### Tools

The genotype-based deconvolution workflow includes five deconvolution methods: Demuxlet^17^, Freemuxlet^26^, Vireo^21^, scSplit ^19^, and Souporcell^20^.

The hashing-based deconvolution workflow includes eight hashing deconvolution algorithms: HTODemux^27^, Multiseq^6^, HashedDrops^16^, Demuxem^10^, gmm-demux^28^, BFF ^23^, demuxmix^29^ and Hashsolo^18^, and one doublet-detection method (Solo^18^).

In addition to the two multiplexing workflows, the hadge pipeline includes a doublet detection method, Solo, which is based on a semi-supervised deep learning approach. Since Solo only identifies singlets without revealing the true donor identity of the droplets, we only use it as a supplementary method.

As genotype-based deconvolution techniques rely on SNPs to distinguish samples in the pools, the workflow also includes a preprocessing component with samtools, Freebayes^35^ and cellsnp-lite^36^ as two separate processes for variant calling. The Freebayes process is designed as per the instruction of scSplit (https://github.com/jon-xu/scSplit) to find variants in pooled samples. To optimize runtime, the process is carried out separately for each chromosome. With an additional filtering step, SNPs with a minimum allele count of 2, minimum base quality of 1 and quality scores greater than 30 from each chromosome are retained and merged. As suggested by Vireo, the Mode 1a of cellsnp-lite is called in the cellsnp-lite process to genotype single cells against candidate SNPs. Two allele counts matrices for each given SNP are generated, one for the reference and another one for the alternative allele, which can be then fed into Vireo.

The hashing-based deconvolution workflow also has a pre-processing step to prepare the input data for both HTODemux and Multiseq based on the Seurat vignette (https://satijalab.org/seurat/articles/hashing_vignette.html). A Seurat object is initialized with the cell containing barcodes for the RNA matrix and HTO raw count matrix. Only the cell barcodes that are in the intersection between RNA and HTO counts are retained. The HTO data is added as an independent assay and normalized using centered log-ratio transformation (CLR).

#### Structure

The hadge pipeline features three distinct modes: *genetic, hashing* and *rescue* mode. In the genetic or hashing mode, the pipeline runs either of the two workflows independently. Following deconvolution in each workflow, the output files are passed to the summary process to generate summary files. Within this module, three CSV files are produced per tool as output, with each column representing a trial conducted during a single run of the pipeline. These output files provide a comprehensive summary of three aspects, including the specified parameters for each trial, the classification of individual droplets as singlets, doublets, or negative droplets, and the assignment of cell barcodes to their respective donors. As multiple tools are executed within a single run, additional CSV files are generated to merge the classification and assignment results from different tools into unified data frames.

In the rescue mode, hashing and genotype-based deconvolution workflows work jointly with the aim (i) to recover the droplets where the classification is discordant between the two approaches and (ii) optionally to extract donor-specific variants from the droplets with coherent classification and to reconstruct donor genotypes for mixed samples, which can then be used to rerun genotyped-based deconvolution as a sanity check prove whether the result is reliable. The pipeline first runs the two workflows in parallel and saves the results of all methods in a single CSV file. Next, the file is passed to the ‘donor matching’ process to associate an identity to the anonymous donors using the droplets where the concordance between one genetic donor and one hashtag is maximized.

The process converts the assignment of two tools into a matrix of binary values, with rows representing cell barcodes and columns representing donors or hashtags. The value is set to 1 if the cell is assigned to the donor or hashtag, and 0 otherwise. The similarity between two matrices is calculated column-by-column using Pearson correlation, and hashtags and donors are matched if they have the highest mutual Pearson correlations. If every donor is paired with a hashtag, the pipeline generates a new assignment of the tools with mapped donors and a heat map to visualize the correlation between the donors and hashtags. If Vireo is the optimal genotype-based deconvolution method in donor matching, the process has the option to extract informative variants from donor genotypes estimated by Vireo. Using the input of cellsnp-lite, genotyped SNPs are first filtered based on the SNPs (read depth > 10) among cells with consistent assignment between Vireo and the hashing tool with which it is compared. Only variants with an overrepresented allele are retained, i.e., the frequency of the alternative or reference allele in the group of cells must be greater than a specified threshold (frequency > 90%). The pipeline compares the genotypes of these variants in cells that have been inconsistently deconvolved and keeps only the SNPs that have the same overrepresented allele in cells with and without consistent assignment. These are candidate variants used to distinguish cells from different donors. The process is performed separately on cells from different donors to retrieve donor-specific informative variants. Finally, BCFtools sorts and indexes the donor genotype from Vireo and filters the donor-specific variants. The samples are renamed by the matching hashtags.

#### Demuxlet/Freemuxlet

Dsc-pileup, Demuxlet and Freemuxlet implemented in popscle (v0.1) were performed one after another. Using the BAM file and filtered barcodes file produced by cellranger^37^ as input, dsc-pileup aggregated reads around common SNPs in the human population, which in the case of Freemuxlet are derived from the 1000 Genomes Project and filtered by cellsnp-lite with minor allele frequency (MAF) > 0.05 as reference variant sites (https://sourceforge.net/projects/cellsnp/files/SNPlist/). Demuxlet/Freemuxlet then uses the pileup files from dsc-pileup to deconvolve cells. We ran these methods in default mode.

#### Vireo

Cell genotypes were generated at common SNPs from the 1000 Genomes Project (https://sourceforge.net/projects/cellsnp/files/SNPlist/) using cellsnp-lite (v1.2.2) with default parameters before performing Vireo. Subsequently, the output of cellsnp-lite was processed by Vireosnp (v0.5.6) to perform the deconvolution with default parameters.

#### Souporcell

Souporcell (v2.0) was run on the BAM file and filtered barcodes file produced by cellranger and the human reference (http://cf.10xgenomics.com/supp/cell-exp/refdata-cellranger-GRCh38-3.0.0.tar.gz). We also used common SNPs from the 1000 Genomes Project^11^ with a minor allele frequency of 2% (provided by https://github.com/wheaton5/souporcell) as input to skip repeated and memory intensive steps, remapping and variant-calling.

#### scSplit

scSplit was executed only after the pre-processing and variant calling modules were completed. The input BAM file was pre-processed by SAMtools (v1.15.1) and UMI-tools (v1.1.2). In the variant calling module, freebayes (v1.2) was performed on the pre-processed BAM file to call variants from mixed samples. Taking the pre-processed BAM file and called variants, scSplit (v1.0.8.2) deconvolved the cell mixture in three steps. The count command of scSplit constructed two count matrices for the reference and alternative alleles. To increase the accuracy of donor identification, a list of common SNPs provided by scSplit (https://melbourne.figshare.com/articles/dataset/Common_SNVS_hg38/17032163) was used to filter the count matrices. The run command identified then cells in the pool to the clusters according to the allele matrices, with doublets being assigned to a separate cluster. Finally, the genotype command predicted individual genotypes for every cluster.

#### HTODemux

HTODemux begins with loading the Seurat object, which was created during the pre-processing module using the Seurat R package (v4.3.0). HTODemux (also included in Seurat R package v4.3.0) was then called with default parameters to deconvolve cells based on clr-normalized HTO counts.

#### Multiseq

Similar to HTODemux, MULTIseqDemux (included in Seurat R package v4.3.0) function was performed on the pre-processed Seurat object, with default parameters allowing for automatic determination of the optimal quantile to use in a range from 0.1 to 0.9 by a step of 0.05.

#### Demuxem

The raw RNA and HTO count data were loaded as a MultimodalData object (pegasuspy Python package v1.7.1). Demuxem then deconvolved cells with at least 100 expressed genes and 100 UMIs in two main steps. The antibody background was first determined based on empty barcodes using the KMeans algorithm. The signal hashtag counts were then calculated using background information, and cells with a minimum signal of 10 were assigned to their signal hashtag.

#### Hashsolo

The process expect start from the raw HTO counts in hdf5 file format into an Anndata object (Scanpy v1.9.1) (solo-sc v1.3). We ran Hashsolo with default parameters, setting the priors for the hypothesis of negative droplets, singlets and doublets each to 1/3.

#### HashedDrops

This process requires as input both RNA and HTO raw couunts. First emptyDrops finds cell-containing droplets, this list of barcodes is then used as input to the HashedDrops call (both algorithms are included in DropletUtils R package v1.18.0). We used HashedDrops in default setting.

#### BFF

BFF accepts raw or preprocessed HTO data, while offering a preprocessing step (ProcessCountMatrix), included in the CellHashR pipeline (CellHashR v.1.14.0). Two different alternatives of BFF are available, “BFF raw” and “BFF cluster”, which apply a different processing on the HTO raw counts. Both methods can be run in parallel and the tool will generate a consensus call between the two. We ran both alternatives for the benchmark.

#### Demuxmix

Demuxmix reads HTO raw counts using Read10X function from Seurat (Seurat R package v4.3.0.1). There are four options available “auto”, “reg”, “regpos” and “naive”. With “auto” the tool can optionally receive a vector with library sizes from the RNA data so as to execute one of the regression mixture models available. In the “naive” mode demuxmix works with the HTO counts alone. We ran demuxmix in naive mode.

#### Gmm-demux

GMM-demux (GMM-demux Python package v.0.2.1.3) expects the HTO raw counts as csv or tsv files and the names of the expected cell hashtags. We ran GMM-demux using tsv files under default parameters.

#### Benchmarking mpxMS-dataset

We were granted early access to a dataset generated in a study of progressive multiple sclerosis (Calliope Dendrou, University of Oxford)^32^. In brief, this dataset includes a multiplexed 3’ single nuclear RNA sequencing dataset of brain tissue from 5 controls and 5 cases of progressive multiple sclerosis (mpxMSdataset). The mpxMS-dataset is divided into two sequencing batches (gx12 and gx38) of 6 donors each, with the individual donors hashed with one of six unique TotalSeq™-A anti-nuclear pore complex antibodies. We obtained the count data generated with Cellranger v3.1.0: 6794833 barcodes and 6794880 barcodes were detected in the raw data of gx12 and gx38, respectively. The number of cells detected in each experiment before deconvolution were 4889 for gx12 and 13184 for gx38.

The pipeline was applied to the mpxMS-dataset in the rescue mode. In the genotyped-based deconvolution workflow, Freemuxlet, Vireo, Souporcell and scSplit were used in the absence of reference donor genotypes. To run the algorithm, the number of samples was set to six. All hashing-based deconvolution methods were called to deconvolute the data. All output files were gathered and passed to the corresponding summary component (R v4.2.2). The results of Vireo and Multiseq were used to map donor identities to hashtags in the donor matching component. Donor genotypes estimated by Vireo were then processed by BCFtools (v1.8). The donor-specific variants were extracted from the donor genotypes, where the cell-variants were filtered by a minimal cell count of 10 and the overrepresented allele at a given SNP was then determined by a 90% cut-off.

Data analysis was performed with scanpy (v1.9.3) and scrublet (v0.2.3). Plots were generated with scanpy (v1.9.3), seaborn (v0.12.2) and matplotlib (v3.7.1).

We generally followed the recommendations given by the developers of the package (https://scanpy.readthedocs.io/en/stable/index.html) and have in part adjusted for this dataset and in accordance with analysis best practices^38^.

For analysis, log-transformation and normalization were achieved with scanpy’s log1p() and normalize_total() function. After this, 50 PCs were generated by principal component analysis (PCA) and dimensionality reduction by UMAP was performed using scanpy’s pca() and umap() functions respectively. Cluster identification was performed using the Leiden algorithm and differential expression of the different clusters was generated using scanpy’s rank_genes_groups() function.

Time benchmarks

We benchmarked the performance of hadge against demuxafy and cellhashR using four samples with different cell numbers. Each pipeline is developed in different frameworks and required different configuration. In the demuxafy pipeline, genotype-based deconvolution methods were called sequentially within the singularity container. The benchmark was run on the mpxMS-dataset batch gx12 and gx38, as well as a reduced test dataset provided by demuxafy, using the same parameters as hadge. Since demuxafy doesn’t provide preprocessing functions, we used hadge’s preprocessing module (which includes *freebayes, samtools* and *cellsnp*) to provide the same input data to hadge-genetic and demuxafy so the benchmarking starts from the same inputs. For the cellhashR pipeline we created a conda environment with all the required dependencies as described in the cellhashr github ^39^. In the hadge-hashing pipeline, each deconvolution method was called in its own conda environment separately. For each pipeline run we allowed 160GB RAM memory and 32 CPU cores.

## Code and data availability

The hadge source code is available at https://github.com/theislab/hadge under the MIT license. Further documentation, tutorials and examples are available at https://hadge.readthedocs.io/en/latest.

Jupyter notebooks to reproduce our analysis and figures including Conda environments that specify all versions are available at https://github.com/theislab/hadge-reproducibility.

The mpsMS-dataset applied in this study is an unpublished dataset obtained directly from the authors ^32^.

## Acknowledgements

We thank Dr Luke Zappia for constructive discussion on the design of the pipeline and the overarching project. We’re grateful for Lisa Sikkema’s input on the figure design. A subset of figure panels were created using Biorender. HBS acknowledges support by the German Center for Lung Research (DZL), the Helmholtz Association (CoViPa - lessons to get prepared for future pandemics), the European Union’s Horizon 2020 research and innovation program (grant agreement 874656 - project discovair) and the Chan Zuckerberg Initiative (CZF2019-002438, project Lung Atlas 1.0). MG-P was supported by the Interdisciplinary Bioscience DTP (BBSRC). CAD was supported by the Wellcome Trust and Royal Society (204290/Z/16/Z). Tissue samples and associated clinical and neuropathological data were supplied by the Multiple Sclerosis Society Tissue Bank, funded by the Multiple Sclerosis Society of Great Britain and Northern Ireland, registered charity 207495.

## Author contributions

FC, XW, and LH contributed equally and have the right to name themselves first in their CV. FC and LH conceived the study. FC, XW, MG, LH implemented the hadge pipeline. XW, MG, FC conducted the benchmarking of the tools. FC, XW, LeH conducted the analysis of the data. HBS, FJT supervised the work. All authors read, corrected and approved the final manuscript.

## Conflicts of interest

L.H. is an employee of Lamin Labs. F.J.T. consults for Immunai Inc., Singularity Bio B.V., CytoReason Ltd, and Omniscope Ltd, and has ownership interest in Dermagnostix GmbH and Cellarity.

## Supplementary Material

**Supplementary Figure 1.**
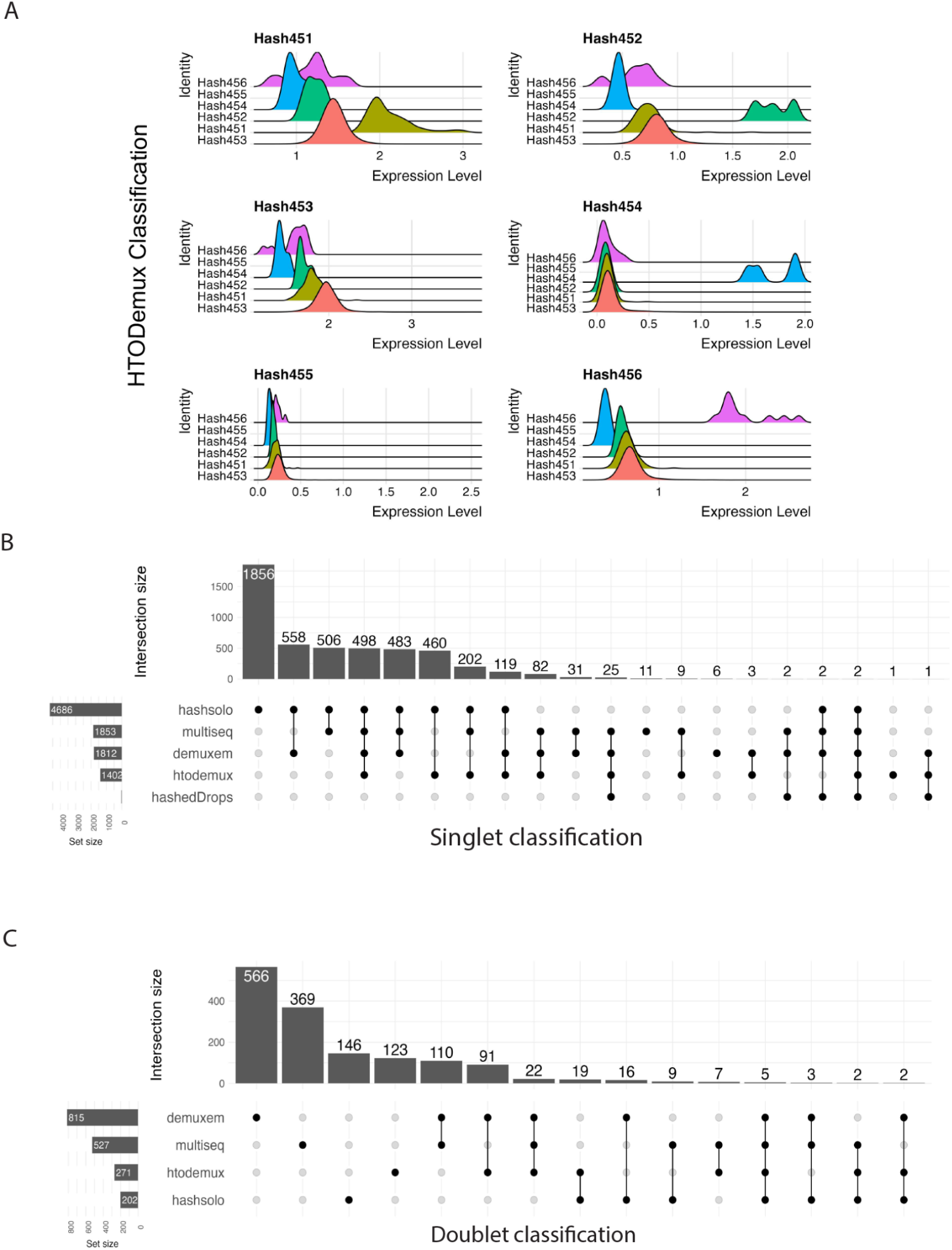
Different performance of hashing-based deconvolution methods. (A) Ridge plot of HTO expression level grouped by cells assigned to different hashtags indicates a relatively similar expression level of different HTOs in cells of Hashtag 454 and 455. (B) Upset plot representing different singlet classification by hashing-based deconvolution methods. Horizontal bars represent the total number of singlets classified by each method. The vertical bars depict the overlapping singlet classifications, indicated by black circles, where singlets are classified by a single method or a combination of methods. (C) Upset plot representing different doublet classification by hashing-based deconvolution methods.

**Supplementary Figure 2.**
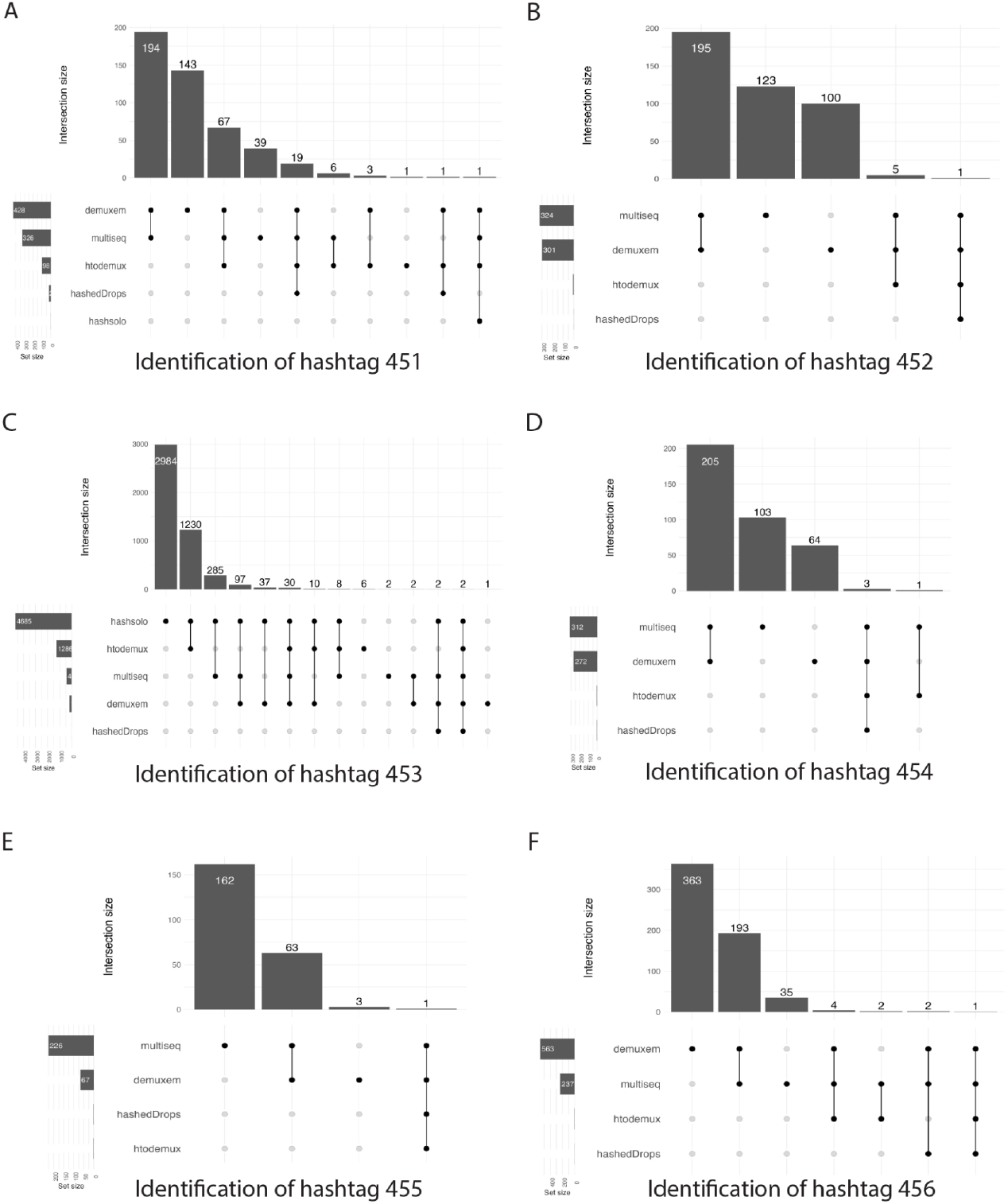
Discordant donor identification performance of hashing-based deconvolution methods on mpxMS-dataset batch gx12.(A) - (E) Upset plot representing different donor assignment of hashing-based deconvolution methods for (A) Hashtag 451, (B) Hashtag 452, (C) Hashtag 453, (D) Hashtag 454, (E) Hashtag 455 and (F) Hashtag 456. Horizontal bars represent the total number of cells assigned to the hashtag by each method. The vertical bars depict the overlapping donor assignment, indicated by black circles, where cells are assigned by a single method or a combination of methods.

**Supplementary Figure 3.**
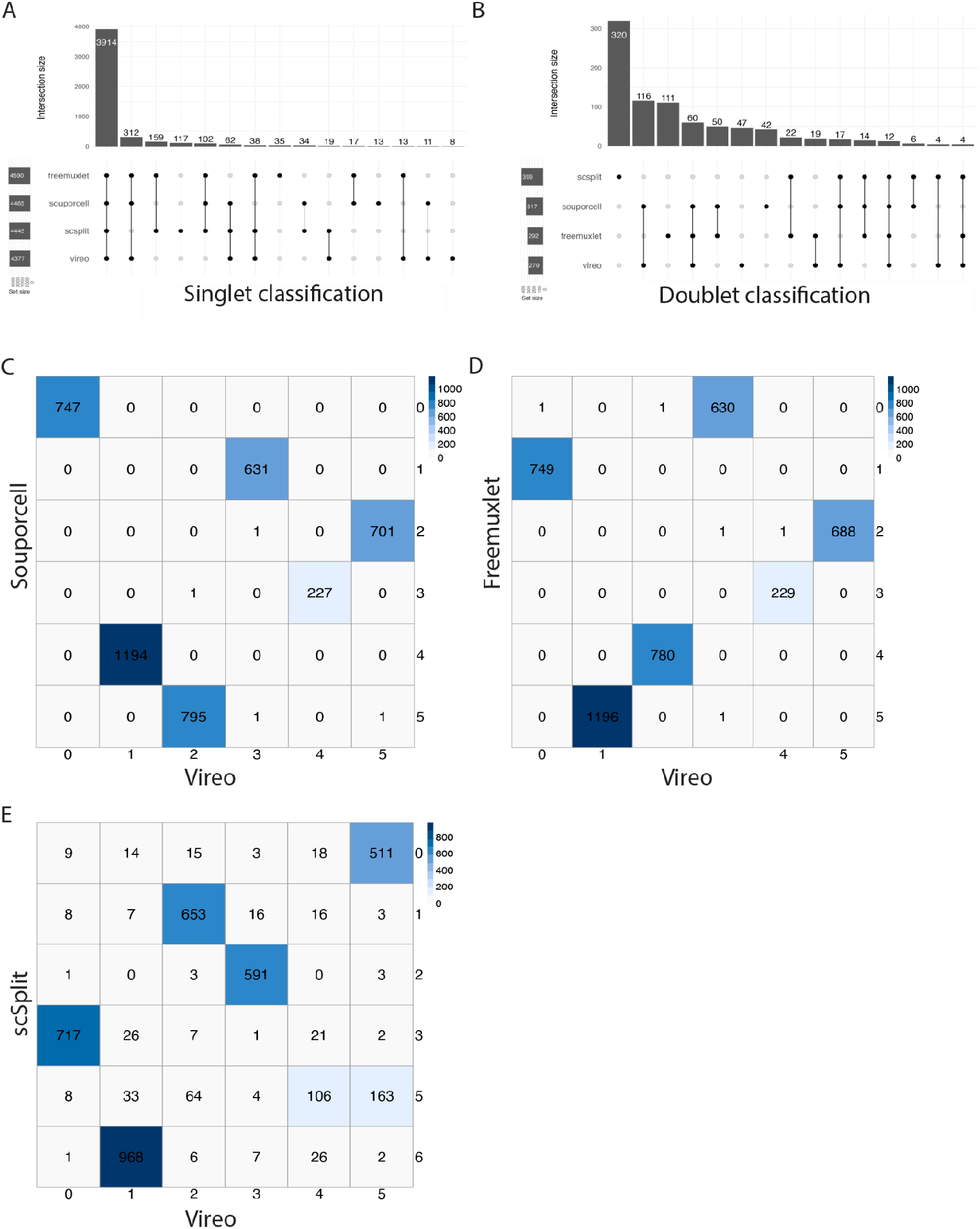
Concordant donor identification performance of genotype-based deconvolution methods on mpxMS-dataset batch gx12. (A) Upset plot representing different singlet classification by genotype-based deconvolution methods. Horizontal bars represent the total number of singlets classified by each method. The vertical bars depict the overlapping singlet classifications, indicated by black circles, where singlets are classified by a single method or a combination of methods. (B) Upset plot representing different doublet classification by genotype-based deconvolution methods. (C) - (E) Confusion matrix representing the agreement in donor identification between Vireo and other three methods: (C) Souporcell, (D) Freemuxlet and (E) scSplit. Vireo is fixed on the x-axis as baseline. The rows and columns represent the anonymous donor clusters genotyped by each respective method. The values within the cells represent the number of singlets assigned to each specific donor cluster.

**Supplementary Figure 4.**
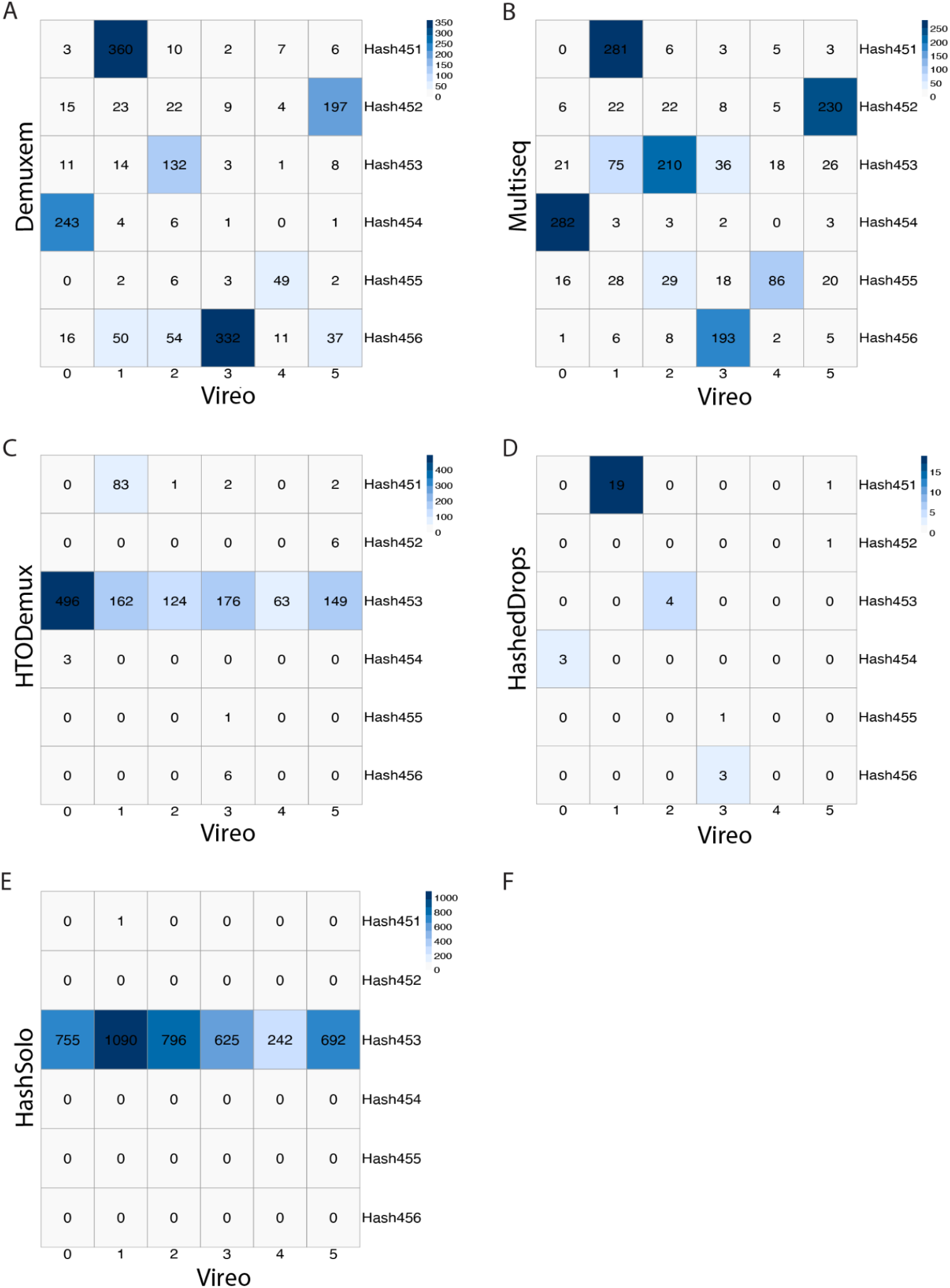
Different donor identification performance between hashing and genotyped-based deconvolution methods on mpxMS-dataset batch gx12. (A) - (E) Confusion matrix representing the agreement in donor identification between Vireo and other five hashing-based deconvolution methods: (A) Demuxem, (B) Multiseq, (C) HTODemux, (D) HashedDrops and (E) HashSolo. Vireo is fixed on the x-axis as baseline. The rows represent the hashtags, while the columns represent the anonymous donor clusters genotyped by Vireo. The values within the cells represent the number of singlets assigned to each specific donor or hashtag.

**Supplementary Figure 5.**
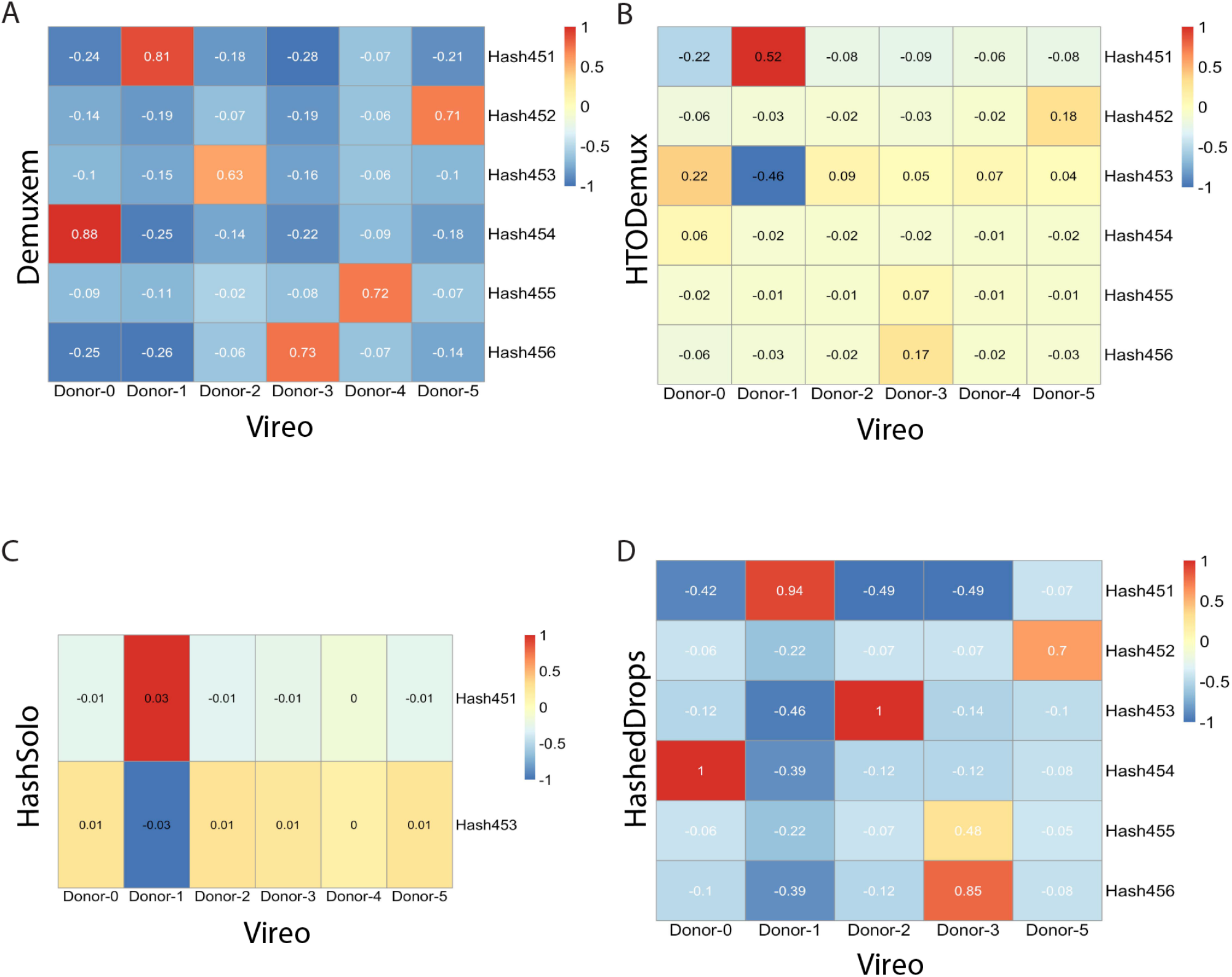
Different donor identification performance between hashing and genotyped-based deconvolution methods on mpxMS-dataset batch gx12 based on Pearson correlation. (A) - (E) Correlation matrix representing the agreement in donor identification between Vireo and other five hashing-based deconvolution methods: (A) Demuxem, (B) HTODemux, (C) HashSolo and (D) HashedDrops. Vireo is fixed on the x-axis as baseline. The rows represent the hashtags, while the columns represent the anonymous donor clusters genotyped by Vireo. The values within the cells represent the Pearson correlation score between singlets assigned to a specific hashtag and those assigned to a donor cluster.

**Supplementary Figure 6.**
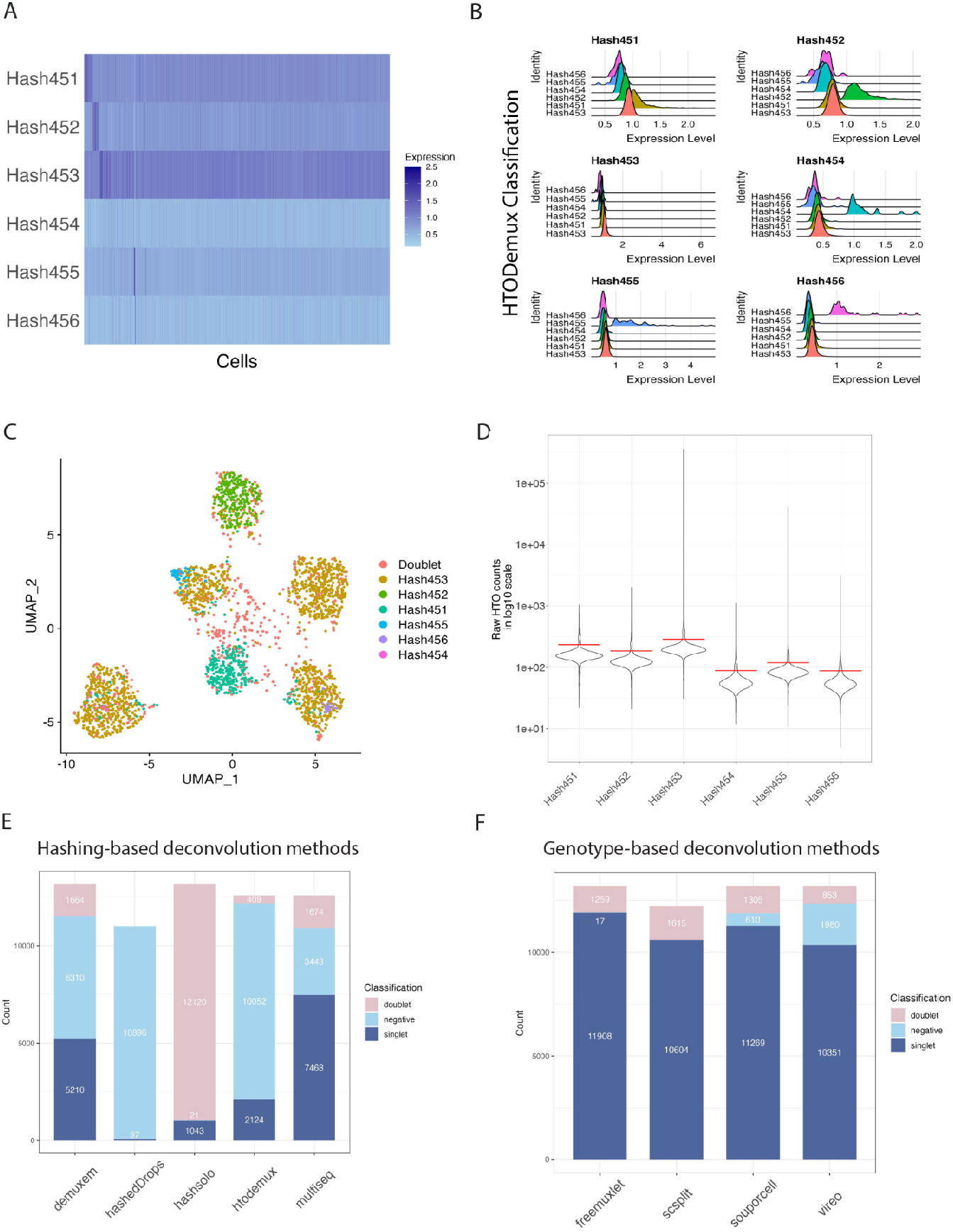
Comparison of the performance of donor deconvolution methods on mpxMS-dataset batch gx38. (A) The heatmap of normalized HTO counts per cell is dominated by Hashtag 453 with noisy or undetectable expression of the other HTOs. (B) Ridge plot of HTO expression level grouped by cells assigned to different hashtags indicates a relatively similar expression level of different HTOs in cells of Hashtag 451 and 453. (C) t-SNE plot of normalized HTO counts colored by HTODemux assignment shows poor separation of the cells based on hashtags, with most droplets assigned to Hashtag 453. (D) The violin plot of raw HTO counts shows a high counts levels of Hashtag 453 in cells compared with the expression of the other HTOs. The (E) Bar plot showing the inconsistent classification of cells by hashing-based deconvolution methods. (F) Bar plot showing a more consistent assignment of the cell mixture to singlets, doublets and negatives by genotype-based deconvolution method.

**Supplementary Figure 7.**
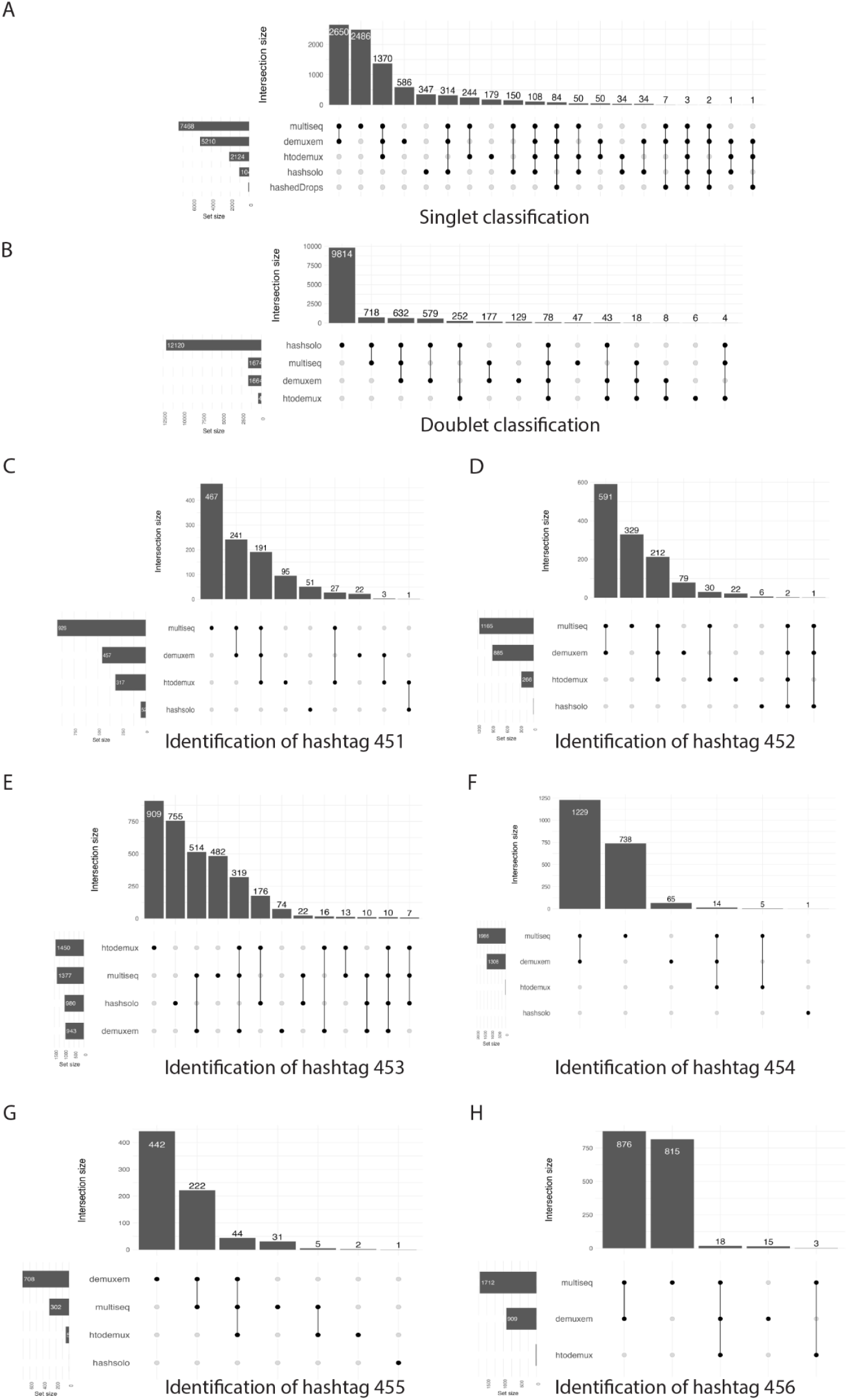
Discordant performance of hashing-based deconvolution methods on mpxMS-dataset batch gx38. (A) Upset plot representing different singlet classification by hashing-based deconvolution methods. Horizontal bars represent the total number of singlets classified by each method. The vertical bars depict the overlapping singlet classifications, indicated by black circles, where singlets are classified by a single method or a combination of methods. (B) Upset plot representing different doublet classification by hashing-based deconvolution methods. (C) - (H) Upset plot representing different donor identification assignment by hashing-based deconvolution methods for (C) Hashtag 451, (D)Hashtag 452, (E) Hashtag 453, (F) Hashtag 454, (G) Hashtag 455 and (H) Hashtag 456.

**Supplementary Figure 8.**
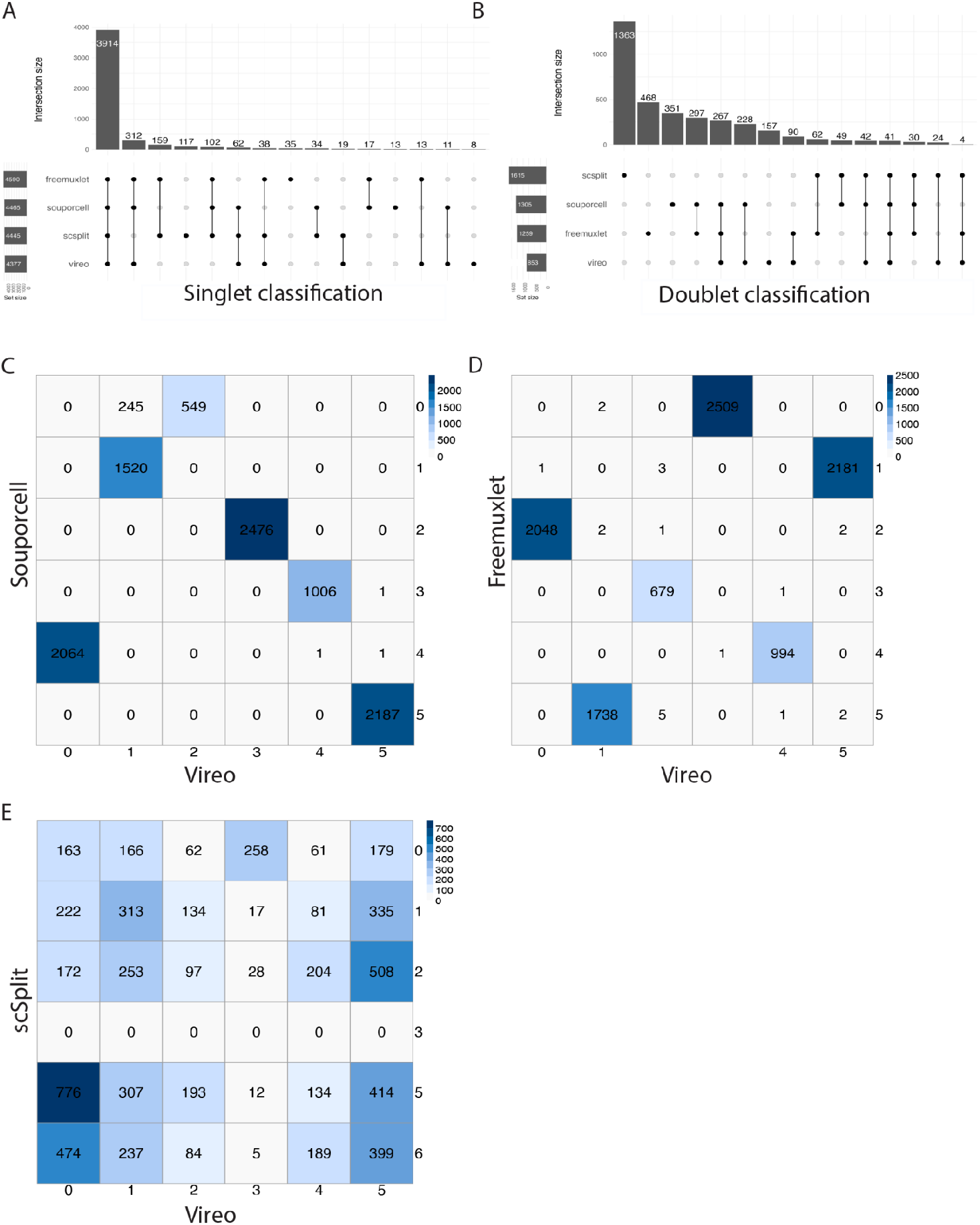
Concordant performance among genotype-based deconvolution methods on mpxMS-dataset batch gx38. (A) Upset plot representing different singlet classification by genotype-based deconvolution methods. Horizontal bars represent the total number of singlets classified by each method. The vertical bars depict the overlapping singlet classifications, indicated by black circles, where singlets are classified by a single method or a combination of methods. (B) Upset plot representing different doublet classification by genotype-based deconvolution methods. (C)-(E) Confusion matrix representing the agreement in donor identification between Vireo and other three methods: (C) Souporcell, (D) Freemuxlet and (E) scSplit. Vireo is fixed on the x-axis as baseline. The rows and columns represent the anonymous donor clusters genotyped by each respective method. The values within the cells represent the number of singlets assigned to each specific donor cluster.

**Supplementary Figure 9.**
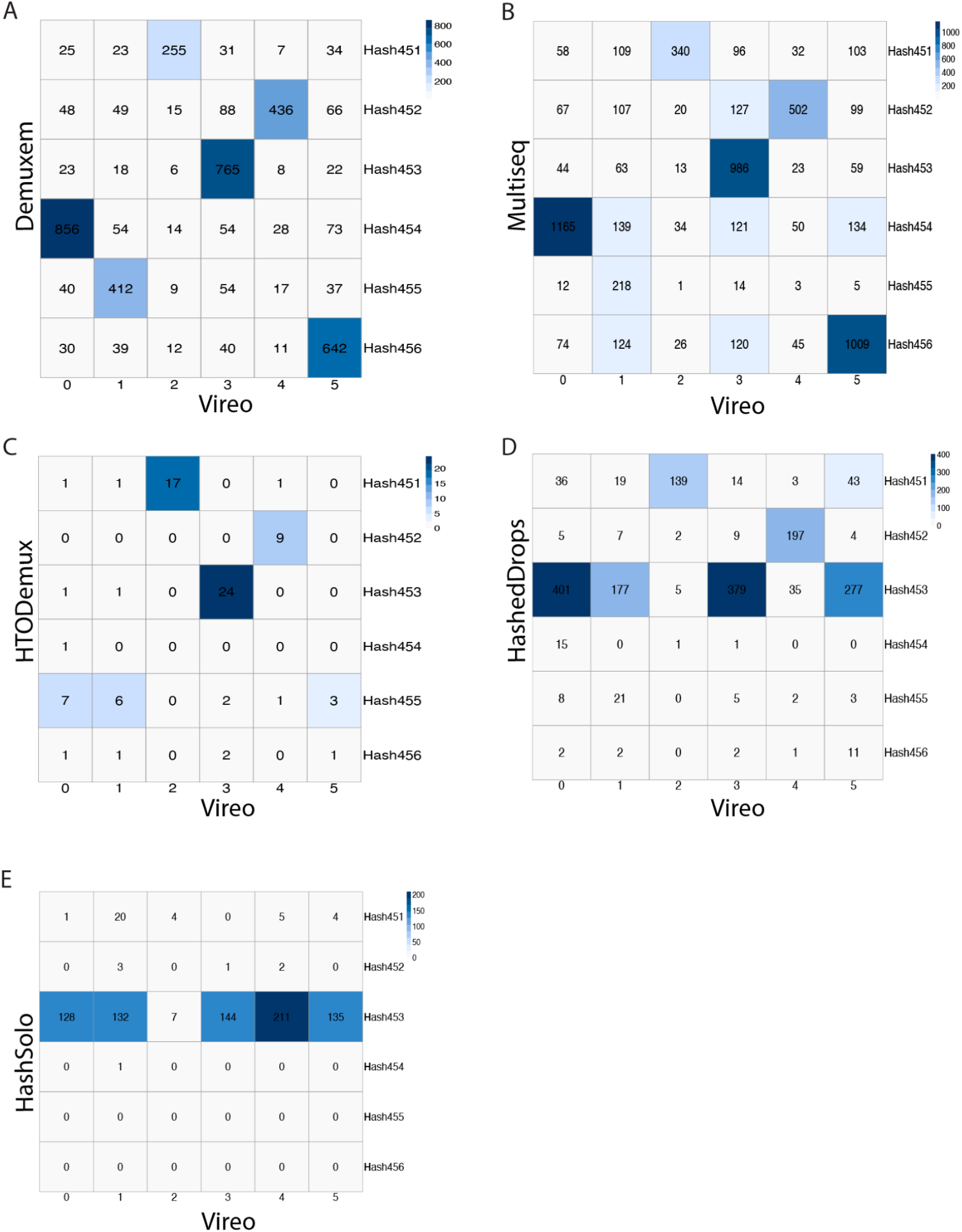
Different donor identification performance between hashing and genotyped-based deconvolution methods on mpxMS-dataset batch gx38. (A) - (E) Confusion matrix representing the agreement in donor identification between Vireo and other five hashing-based deconvolution methods: (A) Demuxem, (B) Multiseq, (C) HTODemux, (D) HashedDrops and (E) HashSolo. Vireo is fixed on the x-axis as baseline. The rows represent the hashtags, while the columns represent the anonymous donor clusters genotyped by Vireo. The values within the cells represent the number of singlets assigned to each specific donor or hashtag.

**Supplementary Figure 10.**
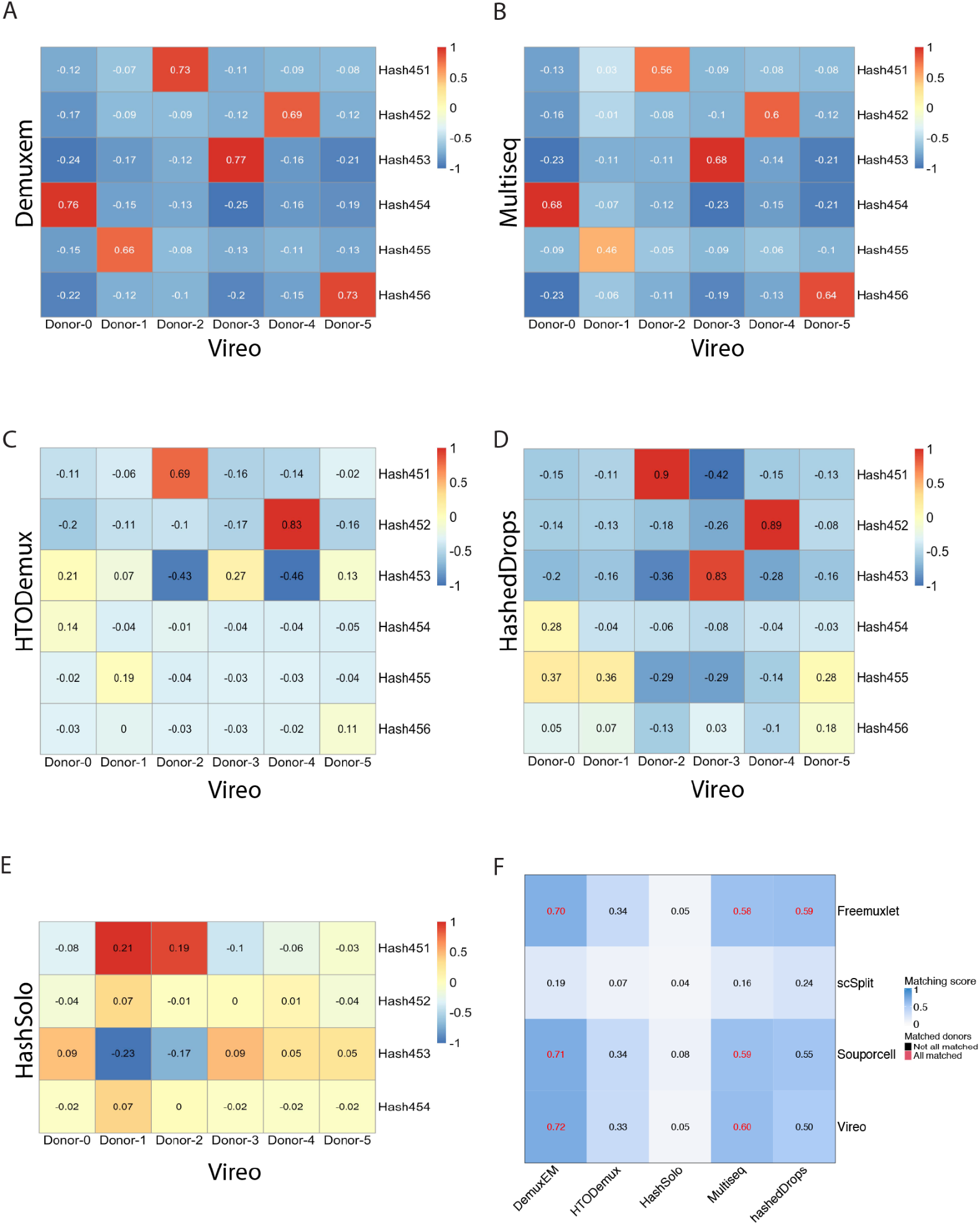
Different donor identification performance between hashing and genotyped-based deconvolution methods on mpxMS-dataset batch gx38. (A) - (E) Correlation matrix representing the agreement in donor identification between Vireo and other five hashing-based deconvolution methods: (A) Demuxem, (B) Multiseq, (B) HTODemux, (D) HashedDrops and (E) HashSolo. Vireo is fixed on the x-axis as baseline. The rows represent the hashtags while the columns represent the anonymous donor clusters genotyped by Vireo. The values within the cells represent the Pearson correlation score between singlets assigned to a specific hashtag and those assigned to a donor cluster. F) Heatmap summarizing the donor matching result shows that DemuxEM and Multiseq are concordant with all genotype-based deconvolution methods except scSplit, where all the donors are matched with a good matching score. The consistency between Freemuxlet and hashedDrops can also be observed.

**Supplementary. Fig. 11:**
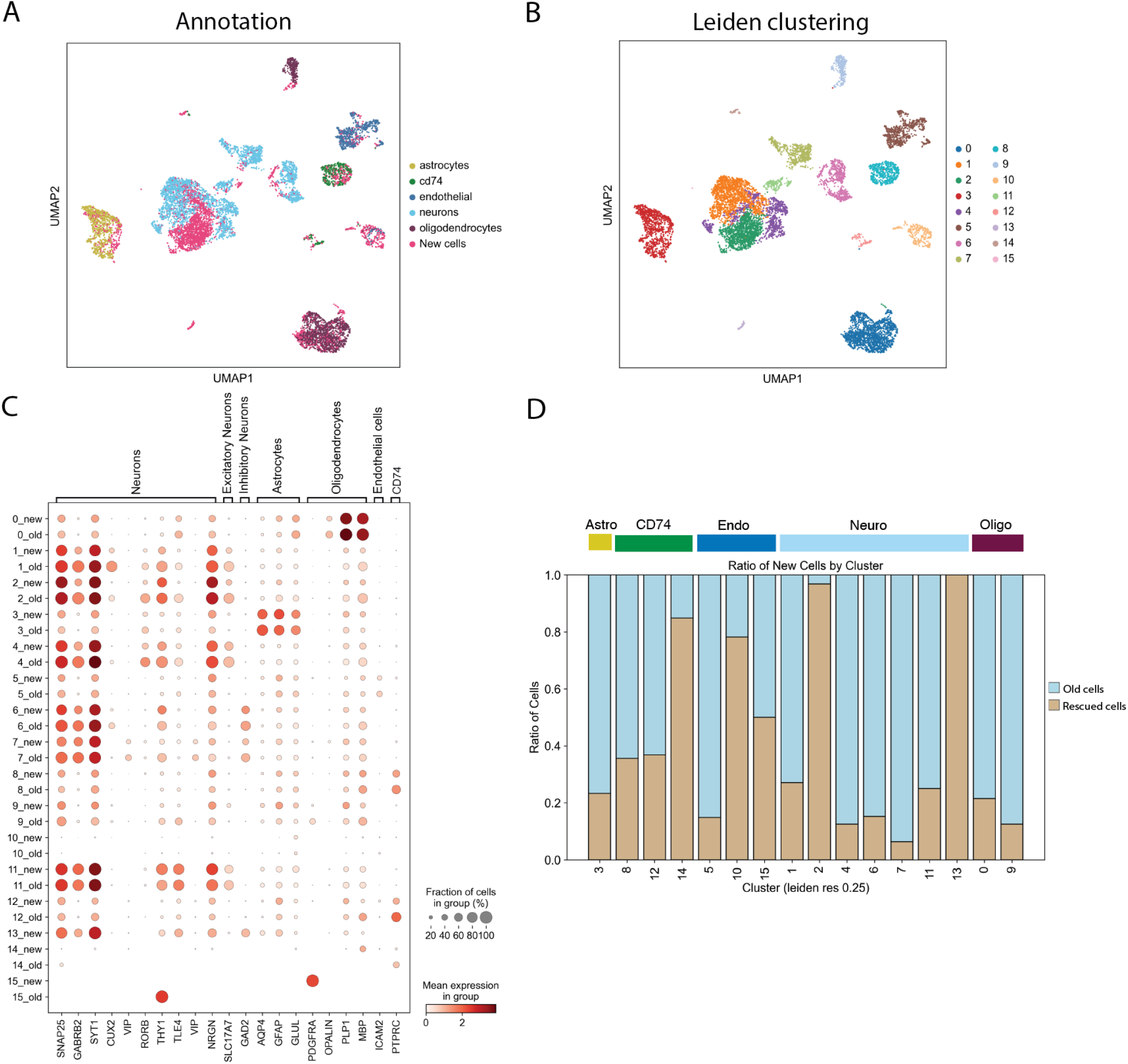
Recovered cells in the second batch (gx38) of the mpxMS-dataset recapitulate original results. (A) UMAP of the single cell gene expression data with old and rescued cells. (B) Leiden clustering of the dataset with old and rescued cells. (C) Dotplot of a selection of marker genes shows concordant expression of markers in old and rescued cells. (D) Barplot showing the cluster-composition in old and rescued cells. Colors on top of barplot identify the cell annotation from (A).

## Notes

### Summary of Updates

small fixes on figures captions

